# Massively parallel single cell lineage tracing using CRISPR/Cas9 induced genetic scars

**DOI:** 10.1101/205971

**Authors:** Bastiaan Spanjaard, Bo Hu, Nina Mitic, Jan Philipp Junker

## Abstract

A key goal of developmental biology is to understand how a single cell transforms into a full-grown organism consisting of many different cell types. Single-cell RNA-sequencing (scRNA-seq) has become a widely-used method due to its ability to identify all cell types in a tissue or organ in a systematic manner ^1–3^. However, a major challenge is to organize the resulting taxonomy of cell types into lineage trees revealing the developmental origin of cells. Here, we present a strategy for simultaneous lineage tracing and transcriptome profiling in thousands of single cells. By combining scRNA-seq with computational analysis of lineage barcodes generated by genome editing of transgenic reporter genes, we reconstruct developmental lineage trees in zebrafish larvae and adult fish. In future analyses, LINNAEUS (LINeage tracing by Nuclease-Activated Editing of Ubiquitous Sequences) can be used as a systematic approach for identifying the lineage origin of novel cell types, or of known cell types under different conditions.

---

Measuring lineage relationships between cell types is crucial for understanding fundamental mechanisms of cell differentiation in development and disease^4,5^. In early development and in adult systems with a constant turnover of cells, short-term lineage predictions can be computed directly on scRNA-seq data by ordering cells along pseudo-temporal trajectories according to transcriptome similarity^6–8^. However, for identifying the developmental origin of cells in the adult body, additional information is required. Genetically encoded fluorescent proteins are widely used as lineage markers^9,10^, but due to limited spectral resolution, optical lineage tracing methods have mostly been restricted to relatively small numbers of cells. Pioneering studies based on viral barcoding^11,12^, transposon integration sites^13^, microsatellite repeats^14^, somatic mutations^15,16^, *Cre*-mediated recombination^17^, and genome editing of reporter constructs^18,19^ have overcome this limitation by using the enormous information capacity of the genome for lineage tracing. However, to date there is no method that allows simultaneous measurement of single-cell transcriptomes and lineage markers *in vivo*.

LINNAEUS is based on the observation that, in the absence of a template for
homologous repair, Cas9 produces short insertions or deletions at its target sites, which are variable in their length and position^18,20,21^ (Fig. 1a). We reasoned that these insertions or deletions (hereafter referred to as genetic “scars”) constitute permanent, heritable cellular barcodes that can be used for lineage analysis and read out by scRNA-seq. To ensure that genetic scarring does not interfere with normal development, we targeted an RFP transgene in the existing zebrafish line *zebrabow M*, which has multiple integrations of the transgenic construct^22^. We injected an sgRNA for RFP and Cas9 protein into 1-cell stage embryos in order to mark individual cells with genetic scars at an early time point in development (Fig. 1b). Loss of RFP fluorescence in injected embryos served as a direct visual confirmation of efficient scar formation (Supplementary Fig. 1). At a later stage, we dissociated the animals into a single cell suspension and analyzed the scars by targeted sequencing of RFP transcripts (see Methods). Simultaneously, we sequenced the transcriptome of the same cells by conventional scRNA-seq using droplet microfluidics^23^ (Fig. 1c and Supplementary Fig. 2).

**Figure 1.**
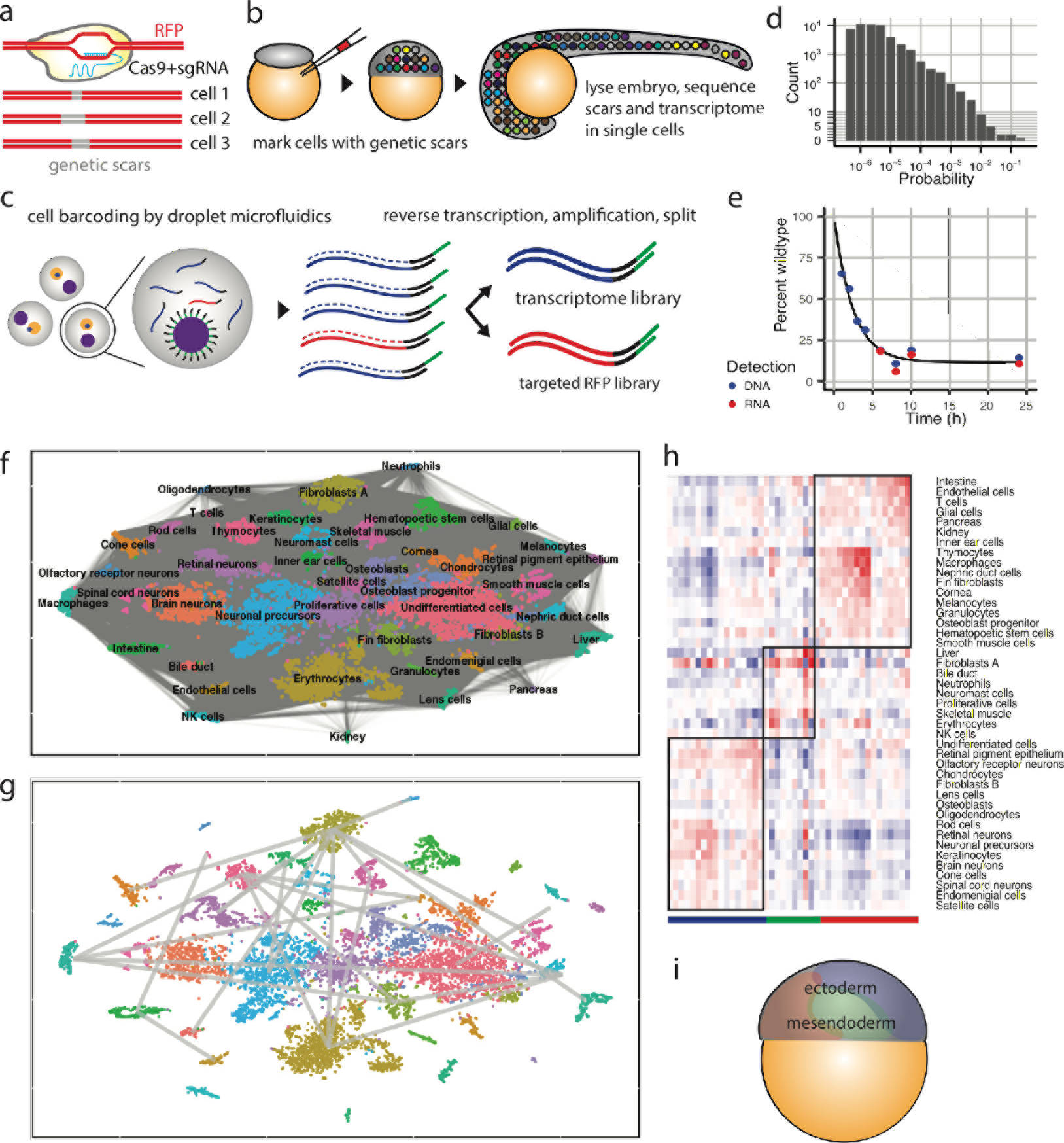
Using the CRISPR/Cas9 system for massively parallel single cell lineage tracing. **(a)** Cas9 creates insertions or deletions in an RFP transgene. These genetic scars can be used as lineage barcodes. **(b)** Sketch of the experimental protocol. Injection of Cas9 and sgRNA for RFP into the zygote marks cells with genetic scars at an early developmental stage. Scars can be read out together with the transcriptome by scRNA-seq at a later stage. **(c)** Approach for simultaneous detection of scars and transcriptome from single cells. Cells are captured by droplet microfluidics, followed by lysis, reverse transcription, and amplification. After amplification, the material is split and processed into a whole transcriptome library and a targeted RFP library for scar detection. **(d**) Probability distribution of scars, measured in bulk experiments. **(e)** Scarring dynamics as measured on the DNA and RNA level, with exponential fit. **(f)** t-SNE representation of scRNA-seq data for dissociated zebrafish larva (5 dpf). Pairs of cells with at least one common scar are linked by gray lines. **(g)** Enrichments of scar connections between cell types compared to random distributions. Only enrichments with p_adj_<0.01 are shown (see Methods). **(h)** Hierarchical clustering of cell types by scar connection strength yields three clusters. **(i)** Cell type clusters determined based on scar connections form contiguous domains on the zebrafish fate map at shield stage. The yolk sac is shown in orange.

We found that Cas9 generated hundreds of unique scars per animal when targeting a single site in RFP (Supplementary Fig. 3), suggesting that analysis of genetic scars constitutes a powerful approach for whole-organism lineage analysis. Bulk analysis of 32 individual larvae revealed that some scar sequences are more likely to be created than others, probably through mechanisms like microhomology-mediated repair^24^ (Fig. 1d). The scars with the highest intrinsic probabilities may be created multiple times per embryo, and are therefore uninformative for lineage reconstruction. We therefore excluded the most frequent scars (p>1%) from further analysis. We found that scarring continued until around 10 hours post fertilization, a stage at which zebrafish already have thousands of cells (Fig. 1e). The early dynamics of scar formation could be described by a simple model using the known cell division rate during early development and a constant per-site scarring rate (Supplementary Fig. 4 and Methods). Thus, our simple injection-based approach for Cas9 induction allowed us to label cells in an important developmental period during which the germ layers are formed and precursor cells for most organs are specified^25^.

Transcriptome analysis of several thousand single cells of dissociated larvae at 5 days post fertilization (dpf) allowed identification of many cell types in the developing zebrafish. Using LINNAEUS, we found around 400,000 links between cells that have shared scars (Fig. 1f). To determine which cell types share common lineage origins, we calculated the enrichment or depletion of scar connections between pairs of cell types compared to random scar distributions. Loss of information due to dropout events is frequent in scRNA-seq, and we noticed that the detected number of scars varies between cell types (from 3 for erythrocytes to 17 for neuromast cells), possibly reflecting differences in cell size or activity of the promoter of the RFP transgene. We therefore designed the background model in such a way that the connectivity distribution of the randomized graphs is identical to the observed graph for all cell types (Supplementary Fig. 5 and Methods). In Fig. 1g we depict lineage connections that are enriched above background with p_adj_<0.01. Clustering of cell types by scar connection strength revealed three groups (Fig. 1h), each of which contains either mostly ectodermal or mesendodermal cell types. These groups formed contiguous domains on the zebrafish fate map^26^, suggesting that they were generated by a small number of early scarring events (Fig. 1i). The clear segregation of scar patterns was reproducible in a second larva (Supplementary Fig. 6), validating our experimental and computational approach.

Next, we set out to analyze the data at higher resolution and reconstruct lineage trees on the single cell level. To do so, we determined the sequence of scar creation events based on our single cell data. Our approach is based on the observation that there is a correspondence between the underlying lineage tree and the resulting scar network graph, a representation of all pairwise combinations of scars that are experimentally observed together in single cells (Fig. 2a). If all scar connections are detected, the scar that is created first has the highest degree of connections in the scar network graph, followed by scars that were created next, enabling lineage tree reconstruction in an iterative manner (Fig. 2b). To account for ambiguity in lineage tree reconstruction caused by missing connections in the scar network graph, we implemented a correction scheme that is based on a maximum likelihood approach. This method evaluates candidate trees by whether or not scar dropout rates are consistent across different branches in a cell type-dependent manner (Supplementary Fig. 7 and Methods). Importantly, our iterative approach for lineage tree reconstruction is robust towards experimental biases that are intrinsic to scRNA-seq data such as scar dropout events. Scar dropouts entail that we do not have full lineage information about every single cell. However, using more than one thousand single cells for tree building allowed us to infer a large part of the missing scar information (Supplementary Fig. 8).

**Figure 2.**
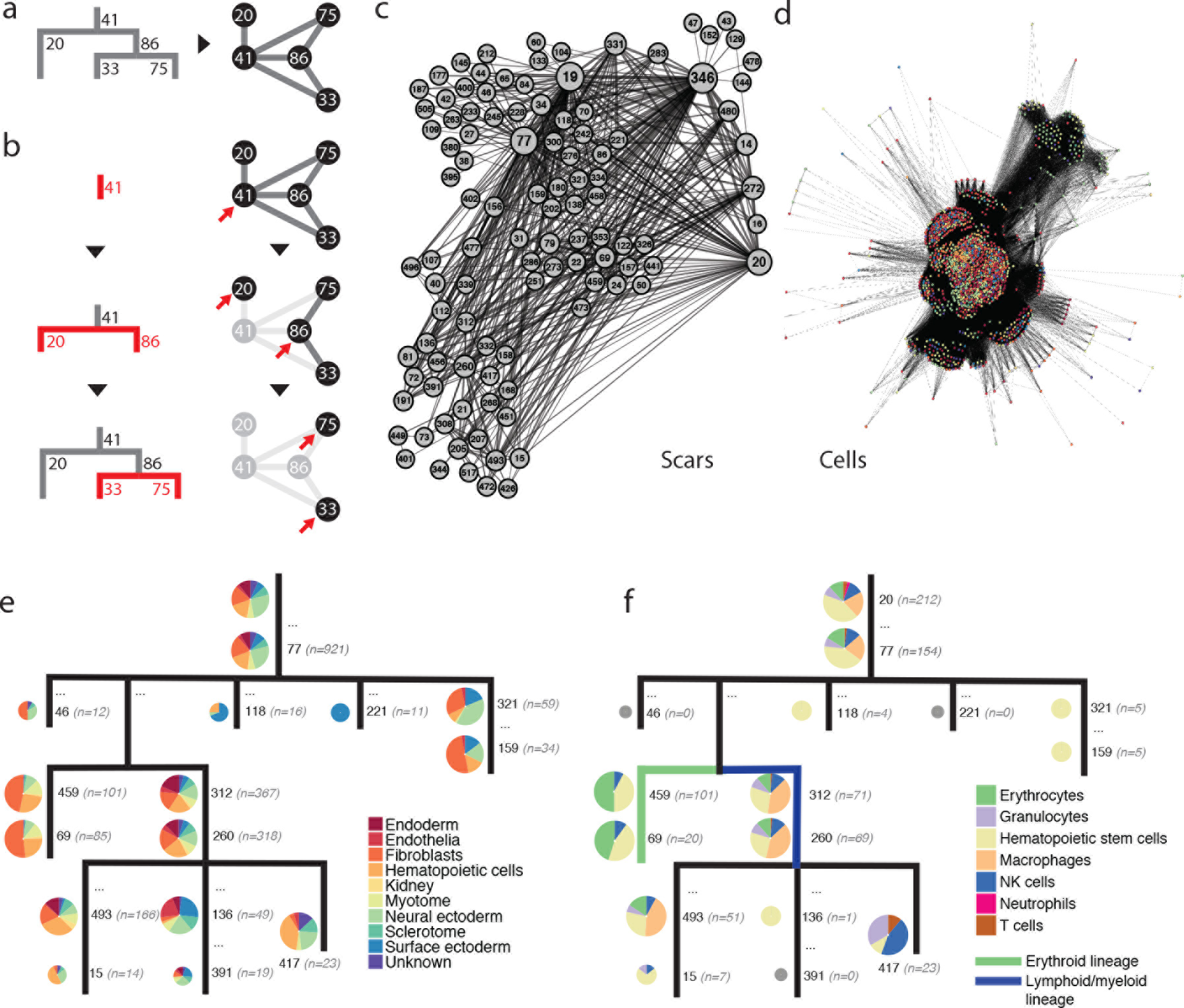
Computational reconstruction of lineage trees on the single cell level. **(a)** Lineage trees can be represented as network graphs. In a scar network graph, each node corresponds to a different scar (designated by a unique scar identification number), and pairs of scars that are co-expressed in single cells are connected by gray lines. **(b)** Cartoon of the computational approach. Network graphs allow reconstructing the order of scar creation events in an iterative approach. The first scar is determined as the one with the highest connectivity (red arrow). Upon removal of the first scar and its connections, the following scars are identified as the most highly connected ones in the reduced network. For details see Methods. **(c)** Scar network graph for 5 dpf larva. Scars are joined if they are co-expressed in at least one cell. Scar identification numbers are determined by the experimentally measured scar probabilities (Fig. 1d), sorted in descending order from high to low probabilities. **(d)** Cell network graph. Cells are joined if they share at least one common scar and are grouped by similarity of scar patterns. Color code indicates tissue of origin as determined by scRNA-seq, see panel (e). **(e)** Lineage tree for 5 dpf larva, including scar identifiers (black font) and cell numbers (gray font). Cell types are grouped into 10 classes as indicated by the color code. Pie charts indicate fractions of cell populations at the individual nodes (cumulative with respect to the branches below). Pie charts are plotted at half size if n<20. **(f)** Lineage tree for 5 dpf larva, zoomed into hematopoietic cell types (see color code).

The scar network graph for the 5 dpf larva exhibited a strong hierarchical structure, with some scars being considerably more highly connected than others (Fig. 2c). A similar structure could be observed in the cell network graph, which also revealed a clear clustering of cell types according to scar profile (Fig. 2d). The reconstructed lineage tree (Fig. 2e, see also replicate experiment in Supplementary Fig. 9) allowed for a more fine-grained analysis compared to connection enrichment analysis (Fig. 1g), revealing for instance that a part of the surface ectoderm splits relatively early from the neural ectoderm. Due to the stochastic nature of cell labeling in LINNAEUS, scar creation is not synchronized with mitosis. It is therefore important to note that reconstructed lineage trees do not necessarily contain all cell divisions. Furthermore, early zebrafish development is highly variable^27^. We can therefore not expect to find exact correspondence of early lineage trees for all cell types in different animals.

For analysis at even higher resolution, we decided to zoom into selected groups of cell types from different germ layers - the hematopoietic system, endodermal cell types, and neuronal cell types. For the hematopoietic system, we find a lineage split between erythrocytes and non-erythroid blood cell types (Fig. 2f). Interestingly, we do not observe complete separation of cell types by lineages. These observations probably reflect the transition from primitive to definitive hematopoiesis in early zebrafish development, as primitive hematopoiesis produces mostly erythrocytes, whereas definitive hematopoietic stem cells are capable of generating all blood cell types ^28^. For endodermal and neuronal cell types, we observed a similar structure of cell-type specific lineage branches, giving rise to different organs, such as the thymus, the hepatopancreas, and the optic apparatus (Supplementary Fig. 10).

During development from embryo to adult, the cell type diversity of tissues and organs increases drastically. One of the major applications of massively parallel single cell lineage tracing will therefore be to systematic identification of the origin of novel cell types. We hence decided to apply LINNAEUS to dissected organs of adult fish in order to explore the resolution of our experimental and computational approach. Analysis of the adult telencephalon, heart, blood, liver, and pancreas by scRNA-seq allowed us to identify many different cell types in these organs (Fig. 3a). Simultaneous measurement of genetic scars enabled us to generate network graphs similar to the ones we obtained for the 5 dpf data. We then determined the order of scar creation events and analyzed the resulting lineage trees at low granularity by grouping the detected cell types into 6 tissue-type categories (brain, endocardium, cardiomyocytes, hematopoietic cells, liver, pancreas). In general, we observed a stronger separation of organs than at 5 dpf (Fig. 3b and Supplementary Fig. 11), which may be caused by inhomogeneous expansion of clones at larval and juvenile stages. We next zoomed into cell types of meso-, endo- and ectodermal origin. Our tree building approach confirmed the split of the three germ layers (Fig 3c, d and Supplemental Fig. 12). Besides identifying numerous clones giving rise to distinct cell types (arrows in Fig. 3c, d), our approach also allowed us to analyze the hierarchy of cell fate decisions, including a separation of alpha/beta and delta/epsilon cells in the pancreas (Fig. 3d) that we also observed in a biological replicate (Supplementary Fig. 11). This is a novel finding, which may be linked to the recent discovery of two waves of endocrine cell differentiation^29^. Interestingly, most neurons and radial glia were found in branches that were fully separate from the main tree, suggesting they are in lineages that split off early (Supplementary Fig. 12). However, the microglia in the longest branch of the tree were placed into a clone that also gave rise to immune cells of the blood.

**Figure 3.**
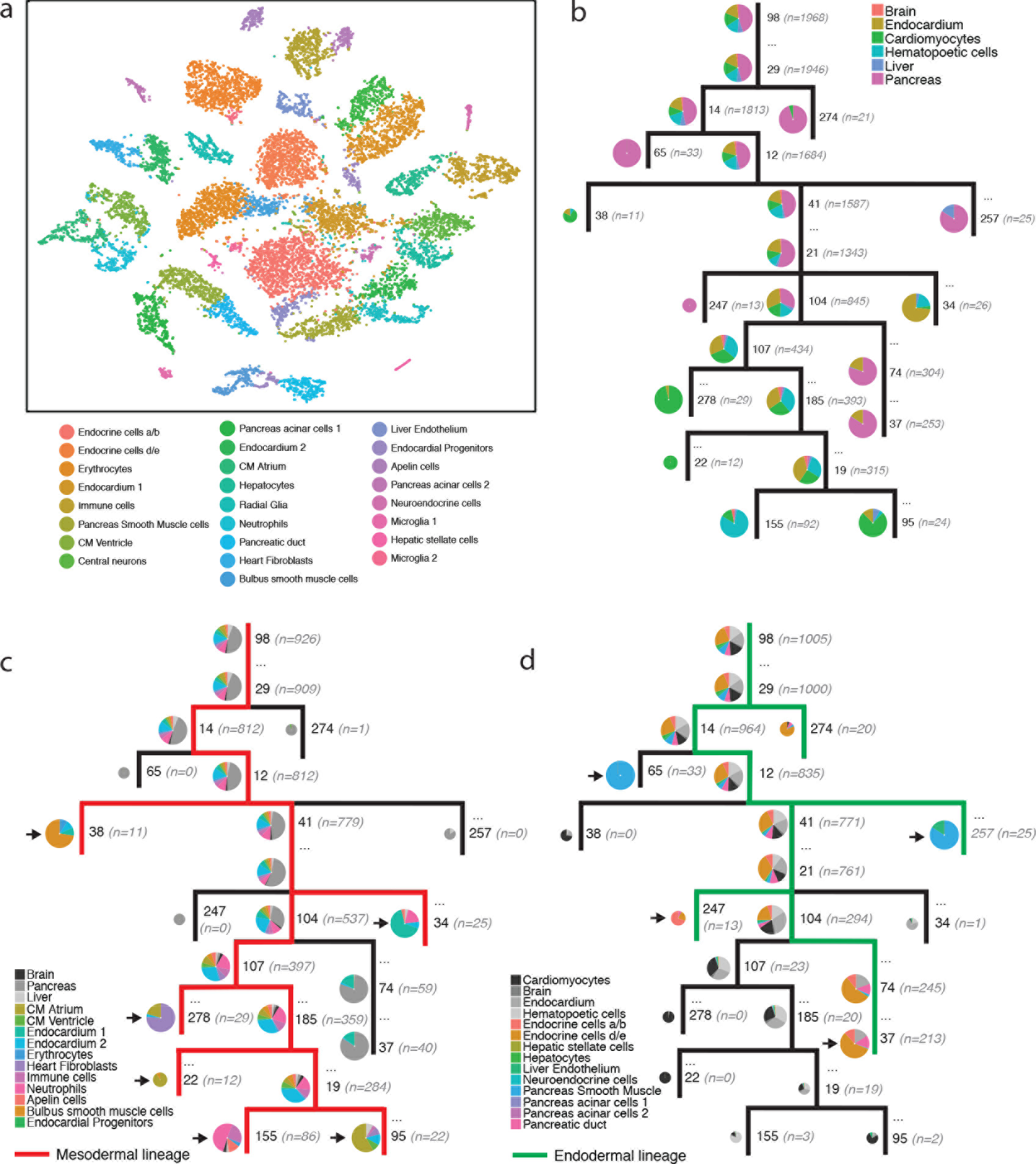
Computational reconstruction of lineage trees on the single cell level. **(a)** t-SNE representation of scRNA-seq data for dissociated organs from adult zebrafish. **(b)** Lineage tree for adult organs, including scar identifiers (black font) and cell numbers (gray font). Cell types are grouped into 5 tissue-type categories. **(c, d)** Fine-grained lineage trees zooming into mesodermal and endodermal lineages. Clones giving rise to distinct cell types are marked with black arrows.

Here, we presented LINNAEUS, a method for simultaneous lineage analysis and transcriptome profiling that is compatible with droplet microfluidics and can be scaled up to thousands of single cells. Importantly, our approach is based on an existing transgenic animal with multiple integrations of a transgenic construct, which should facilitate adaptation of the method to other model systems. An important advantage of our strategy compared to competing technologies such as viral barcoding and other inducible sequence-based lineage tracing methods is the ability to move beyond clonal analysis and to computationally reconstruct full lineage trees on the single cell level. This is made possible by our computational approach for tree reconstruction that is robust to dropout events, and by our experimental strategy that uses independent scarring sites whose scars, once created, cannot be changed again. Within a single experiment, data analysis can be performed at different levels of granularity, from germ layers to organs and cell types. Our combined experimental and computational platform thus provides a powerful strategy for dissecting the lineage origin of uncharacterized cell types and for measuring the capacity of lineage trees to adapt to genetic or environmental perturbations. We anticipate that future modifications of the experimental platform, such as for instance inducible systems, will enable longer periods of lineage tracing, and molecular recording of cellular signaling events.

## Acknowledgements

We thank Robert Opitz, Sharan Janjuha, Nikolay Ninov, Mariana Guedes Simoes and Daniela Panakova for help with cell type identification, Dominic Grün for code to extract single-cell mRNA transcript counts, and we acknowledge support by MDC/BIMSB core facilities (genomics, bioinformatics, zebrafish). This work was funded by a European Research Council Starting Grant (ERC-StG 715361 SPACEVAR). BH was supported by a PhD fellowship from Studienstiftung des deutschen Volkes.

### Author contributions

JPJ, BS and BH conceived and designed the project. BH developed the experimental protocol, and BH and NM performed experiments. BS developed computational methods and analyzed the data. JPJ guided experiments and analysis. JPJ and BS wrote the manuscript, with input from all other authors. All authors discussed and interpreted results.

## Methods

### Zebrafish lines and animal husbandry

We used the transgenic zebrafish line *zebrabow M*^22^ for LINNAEUS. This line has multiple integrations of a transgenic construct that expresses RFP from the ubi promoter, which is constitutively active in all cell types. Fish were maintained according to standard laboratory conditions. Animal experiments were done in compliance with German and Berlin state law, carefully monitored by the local authority for animal protection (Lageso). Embryos of the *zebrabow M* line were injected at the 1-cell stage with 2 nl Cas9 protein (NEB, final concentration 350 ng/μl) in combination with an sgRNA targeting RFP (final concentration 50 ng/μl).

The sgRNA was in vitro transcribed from a template using the MEGAscript^®^ T7 Transcription Kit (Thermo Scientific). The sgRNA template was synthesized with T4 DNA polymerase (New England Biolabs) by partially annealing two single stranded DNA oligonucleotides containing the T7 promotor and the RFP binding sequence, and the tracrRNA sequence, respectively. In the experiments described here, we did not use the ability of the line *zebrabow M* to switch from RFP to YFP or CFP expression upon addition of Cre^22^.

### Scar detection in bulk samples

DNA-based scar detection: DNA of single animals was extracted by heating the samples in 50 μl of 50 mM NaOH at 95°C for 20 minutes. 1/10 volume of 1 M Tris-HCl, pH = 8.4 was then used to neutralize the mixture. We took 20 μl of the DNA for amplification of scar sequences using RFP-specific barcoded primers. The RFP primers were chosen such that the cut site of Cas9 was positioned approximately in the middle of the sequencing read. We then pooled the PCR products, performed a clean-up reaction using magnetic beads (AMPure Beads, Beckman Coulter), and added Illumina sequencing adapters in a second PCR reaction.

RNA-based scar detection: RNA of single or pooled animals was extracted with TRIzol™ Reagent (Thermo Fisher Scientific) according to the manufacturer’s protocol. The RNA was precipitated using isopropanol, and the pellet was washed 2 times with 75% ethanol, air dried, and resuspended in 10 μl of reverse transcription mix (0.3 μM poly-T primer, 1x first strand buffer (Thermo Fisher Scientifc), 10 μM DTT, 1 mM dNTPs, 0.5 μl RNAseOUT™ (Thermo Fisher, Cat. No. 10777019), 0.5 μl SuperScript™ II (Thermo Fisher, Cat. No. 18064-014. The reaction was incubated at 42°C for 2 h for reverse transcription, followed by scar specific PCR amplification as described above for DNA-based scar detection.

### Transcriptome and scar detection in single cells

Single cells were captured using Chromium™ (10X Genomics, PN-120233), a droplet-based scRNA-seq device according to the manufacturer’s recommendations. Briefly, the instrument encapsulates single cells with barcoded beads, followed by cell lysis and reverse transcription in droplets. Reverse transcription was performed with polyT primers containing cell-specific barcodes, Unique Molecular Identifiers^30^ (UMI), and adapter sequences. After pooling and a first round of amplification, the library was split in half. The first half was fragmented and processed into a conventional scRNA-seq library using the manufacturer’s protocols. We used the second, unfragmented, half to amplify scar reads by two rounds of PCR, using two nested forward primers that are specific to RFP, and reverse primers binding to the adapter site. The RFP primers were chosen such that the cut site of Cas9 was positioned approximately in the middle of the sequencing read, ensuring that a broad range of deletion lengths can be reliably detected. We confirmed successful library preparation by Bioanalyzer (DNA HS kit, Agilent). Samples were sequenced on Illumina NextSeq 500 2x 75 bp and Illumina HiSeq 2500 2x 100 bp.

### Mapping and extraction of single cell mRNA transcript counts

Every sequencing read consists of a cellular barcode, a UMI, and a transcript sequence originating from an mRNA molecule. These transcript sequences were aligned using *bwa aln*^31^ with setting ‘-q 50’ to a reference transcriptome constructed from Ensembl release 74 (www.ensembl.org) with extended 3’ UTR regions^32^. We filtered out all unmapped reads and all reads that were not uniquely mapped. After alignment, we determined which cellular barcodes corresponded to cells. We defined a cell to be a cellular barcode with at least five hundred uniquely mapped molecules.

For each cellular barcode, we counted the number of molecules mapped to each gene, using the UMI-correction method described by Grün et al.^33^. This method corrects for the possibility of the same UMI being used for two different transcripts in the same cell with the formula *t* = −*K ln(1 − k_o/K)*, with *t* the final number of transcripts, *k_o* the observed UMIs, and *K* the total number of UMIs possible.

As protection against barcode sequencing errors, we counted the occurrence of each nucleotide for each barcode and filtered out barcodes in which one nucleotide occurred ten or more times. Furthermore, we filtered out barcodes that were one nucleotide substitution removed from a barcode with at least eight times as many transcripts.

### Mapping and filtering of single cell scar data

Scar reads have the same structure as transcript reads: they consist of a barcode, a UMI and a scar. The scar sequences were aligned using *bwa mem*^34^ to a reference of RFP. We defined a cell as a barcode with at least 500 reads. We removed reads that were unmapped, had an incorrect barcode, or did not start with the exact PCR primer we used. We truncated all scar sequences to 75 nucleotides, removed shorter sequences, and filtered the data for sequencing errors and doublets.

### Determination of scar probabilities

We aligned DNA-amplified reads of thirty-two embryos to a reference of RFP. We filtered out unmapped reads and reads that did not start with the exact PCR primer, and truncated all reads to one hundred nucleotides, removing shorter ones. To determine the creation probabilities of the different scars, we removed all unscarred RFP reads from each embryo. We normalized the scar content of each embryo to one and calculated scar probabilities as the average ratio with which each scar was observed.

To account for the slightly different sequencing read structure of single cell and bulk scar detection (see above), we considered only the nucleotides that are shared between the two approaches, and we assigned the bulk scar probabilities to single cell scars accordingly. Since scars with a high creation probability contain little lineage information, we conservatively removed all scars for which we cannot determine the probability in bulk. We filtered out all single cell scars that were not detected in bulk experiments and that were partially soft-clipping, meaning they contained nucleotides that cannot be aligned to RFP or identified as insertions or single nucleotide polymorphisms. Single cell scars that were not detected in bulk but that did not have any soft-clipping nucleotides are indeed detectable, so we set their probability to the lowest probability value detected in bulk.

### Determination of scarring dynamics

Embryos were injected with Cas9 and sgRNA at the 1-cell stage. After 1, 2, 3, 4, 6, 8, 10, and 24 hours, 2-3 embryos were collected and RNA and/or DNA were extracted using TRIzol Reagent according to the manufacturer’s protocols. Bulk scar libraries were produced as described above. For each sample, we calculated the percentage of unscarred RFP. We fit a negative exponential to this data, assuming that the fraction of unscarred RFP at t=0 was one.

### Identifying cell types

We used the R package ‘Seurat’, version 1.4.0.9^35^, for cell-type identification as described below. We removed genes that were not found in at least three cells, and removed cells that had less than two hundred of those genes. We log-normalized the transcript counts and removed cells with more than 2,500 genes observed. For single cells from 5 dpf larvae, we filtered out cells with a mitochondrial content of more than 7.5 percent, and for single cells from adult organs we filtered out cells with a mitochondrial content of more than thirty percent; we expect the cardiomyocytes in particular to have high mitochondrial content. We regressed out influences of the number of transcripts and mitochondrial transcripts, and kept a total of 4071 highly-variable genes for cells of 5 dpf larvae, and 3303 highly-variable genes for adult organ cells. We performed a principal component analysis and kept the first sixty components for single cells from 5 dpf larvae, and twenty-two for adult organ single cells. Clustering, using the smart local moving algorithm^36^ on a K-nearest neighbor graph of cells with resolution 1.8 was done on these components. Dimensional reduction, using t-Stochastic Neighbor Embedding^37,38^ (tSNE), was done on the sixty components for the 5 dpf larvae, and on components three to twenty-two for the adult organs to reduce the visual impact of batch effects. Differential gene expressions were calculated using a likelihood-ratio test^39^ for all clusters, and these were used to identify the cell types. Cells isolated from the adult pancreas, liver or heart but assigned a neuronal identity were removed. Clusters were subsequently merged if they were found to have the same cell type.

### Connection enrichment analysis

We used an analysis of the scars shared between cells to illuminate the overall structure of the sequencing results from 5 dpf larvae. We expect that cells in which we observe the same scar have a shared lineage. To understand the scarring process better, we aimed to find out which cell types share many scars — these cell types would have a strong lineage relationship — and which cell types do not share many scars — these cell types would not have many immediate shared precursors.

We call cells ‘connected’ if they share at least one scar that has a creation probability of less than 1%. All such connections for one animal are shown in Fig 1f. To find out whether cell types have a higher number of connections between them than expected by chance, we developed the background model described below (see also Supplementary Fig. 5). The background model starts with the realization that a connection is defined by its endpoints, and that therefore the number of expected connections between two cell types is determined by the number of connection endpoints of the two cell types. More precisely, the chance of forming a connection between cell type A and B is given by *p(A-B)* = *2* * *CE(A)*\**CE(B)/CE(tot)*^*2*, and that of forming a connection within cell type A by *p(A-A)* = *CE(A)*^*2/CE(tot)*^*2*, with *CE(A)* the number of connection endpoints of cell type A, and *CE(tot)* the total number of connection endpoints. These probabilities define a binomial background model. Using this model, we calculate the enrichment z-score between cell types, i.e. how many standard deviations the observed number of connections between two cell types is away from the expected number of connections. A positive enrichment score indicates more connections than expected by chance, a negative enrichment score indicates less connections than expected by chance.

We define the distance between cell types based on their enrichment z-scores by the following equation: *D(A, B)* = *1* − *(E(A, B)* − *E*_*min*_*)/(E*_*max*_ − *E*_*min*_), with *D(A, B)* the distance between cell types A and B, *E(A, B)* the enrichment z-score between them, *E*_*min*_ the minimal enrichment z-score and *E*_*max*_ the maximum enrichment z-score. The term *E* − *E*_*min*_ can be understood as a translation of all enrichment scores to positive values. These values are then divided by the maximum value and subtracted from 1 to create distances scaled between 0 and 1. We performed hierarchical clustering on these distances, using average linkage as implemented by the *hclust* function in R. We cut the dendrogram into three clusters for larva 1, shown in Fig. 1h, and into two clusters for larva 2, shown in Supplementary Fig. 6.

### Tree building

To build single-cell lineage trees, we start by creating a graph of scars. For this, we considered all scars that have a probability lower than 1% and that we observed in at least two cells. In this graph, scars are represented by nodes and nodes are connected if the scars coincide in two or more cells.

In Fig. 2a we show this graph in a simple example, assuming all scars present are always observed. In this idealized example, a scar created in a cell will coincide with all scars created in the progeny of that cell. This means that in the graph of scars, scars created early will have the most connections. We can use this fact to infer the creation order and branching of scars (afterwards referred to as scar tree) in a recursive manner, starting with an empty scar tree:

1. We select the most-connected scar present in the graph.
2. We remove this scar from the graph and place it at the relevant tip in the scar tree.
3. Underneath this tip, we create as many branches as we have components in the scar graph after removing the scar.
4. We iterate over these graphs, starting from step one above.

For a simple tree, this process is illustrated in Fig. 2b. The above procedure would work for the idealized case where all scars present are always observed. In our actual data, however, due to dropout events we cannot assume this is true. We therefore create trees where the order in which we remove scars is predetermined, instead of being determined by their degree in the scar graph. Once we have created such a scar tree and its corresponding cell lineage tree, we can determine how likely our data is, given the tree, allowing for a maximum likelihood tree building procedure.

To determine the likelihood of a scar tree, we first use it to build a cell lineage tree. Using the scar tree as a skeleton, we place all cells in the lineage tree by determining the positions of their scars in the scar tree. Every cell is placed as low as allowed by their scars. If a cell has conflicting scars, i.e. scars that should not occur together according to the scar tree, it is not placed in the tree. Finally, we collapse lineage tree nodes upwards if they have less than five (for adult organs) or ten (for 5 dpf larvae) cells.

The criterium we use to determine the likelihood of a tree is based on scar dropout rates (see illustration in Supplementary Fig. 7). If, for example, according to the tree scar 20 follows scar 41, all cells in which we observe scar 20 should also have scar 41. If scar 75 also follows scar 41, all cells in which we observe 75 should also have scar 41. We may not always observe scar 41 in these cells. But within the sets of cells where we observe either 20 or 75, the ratios of cells in which we observe 41 should be comparable. This scheme gives rise to our likelihood computation: we test the probability that these ratios are found to be unequal by chance, using a two-tailed Fisher’s exact test. We do this test for every pair of scars, testing the dropout rates for every scar that is higher in the scar tree, for every cell type to account for different dropout rates for different cell types. The product of p-values for all these tests is the likelihood of the tree.

We search the tree space using a Metropolis-Hastings Monte Carlo Markov Chain. For the first tree we build, we place scars in the order by their degree in the scar graph. For consecutive trees, we interchange the position of two, three, four or five scars in the order used for the last tree, with probabilities 50%, 30%, 15% and 5% respectively. We then compare the last created tree with the previous tree and decide whether to accept the last created tree or to reject it and use the previous tree as a basis for further trees. We have two criteria for this comparison: First, we set a maximum tree depth, defined as the highest number of consecutive scarring events implied in a scar tree. Trees that have higher depth than this are automatically rejected, unless the last tree also had a higher depth, in which case the tree with the lowest depth is kept. This scenario can occur because the first tree is built without boundary conditions on depth. Secondly, if both trees have lower than maximum depth, we calculate the likelihood for the tree we built and compare it to the likelihood of the previous tree. If its likelihood is greater, we accept the tree and create a new scar order based on the scar order of the last tree as described above. If its likelihood is smaller, the chance of accepting the tree is the ratio of the new and old likelihood. If a tree is rejected, we create a new scar order based on the scar order of the previous tree. After the Markov chain is terminated, we select the tree with the highest likelihood from all trees with a depth that does not exceed the maximum depth.

For larva 1 (Fig. 2), we followed the above protocol with a maximum depth of 22. We only scar connections established at least two separate cells. To reduce computational complexity in this dataset with very high cell type and scar diversity, we selected a single cell type, Fibroblasts A, for tree likelihood testing. This cell type was used due to a high number of cells and a high average number of scars per cell. We created 17,000 trees and were able to place 1300 out of 1471 cells in the tree shown in Fig 2e. For larva 2, we followed the above protocol with a maximum depth of 16. After 30,000 iterations, we were able to place 994 out of 1052 cells in the tree shown in Supplemental Fig 9. For adult 1, we followed the above protocol with a maximum depth of 10. After 30,000 iterations, we were able to place 1299 out of 1323 cells in the tree shown in Fig. 3b. For adult 2, we followed the above protocol with a maximum depth of 24. After 28,000 iterations, we were able to place 1968 out of 2033 cells in the tree shown in Supplemental Fig 11. For visual simplification, we left out a subset of scars when plotting the trees. For adult 2, we left out any nodes that did not have at least ten cells. Furthermore, on vertical branches we only plotted the top and bottom node. Nodes that are not plotted are indicated by “…”.

### Graphs

Cell and scar graphs were made using Gephi 0.9.1 (https://gephi.org). We used the Fruchterman Reingold algorithm for the layout of the cell graphs, and the Yifan Hu algorithm for the layout of the scar graphs. The scar graphs were afterwards edited manually.

### Simulations

We simulated the scarring process during embryo development (Supplementary Fig. 4). To do this, we used a simple model that starts with one cell, and in which all cells present undergo synchronized mitosis (using cell division rates measured by microscopy^40,41^). Every cell cycle, the RFP integrations of the cells can acquire a scar.

The chance of creating a scar is fixed for every integration for every cell division, and scars are transmitted to a cell’s progeny. A scarring rate of 0.3 per hour reproduced the fit scarring dynamics during the first three hours.

---

**Supplementary Figure 1.**
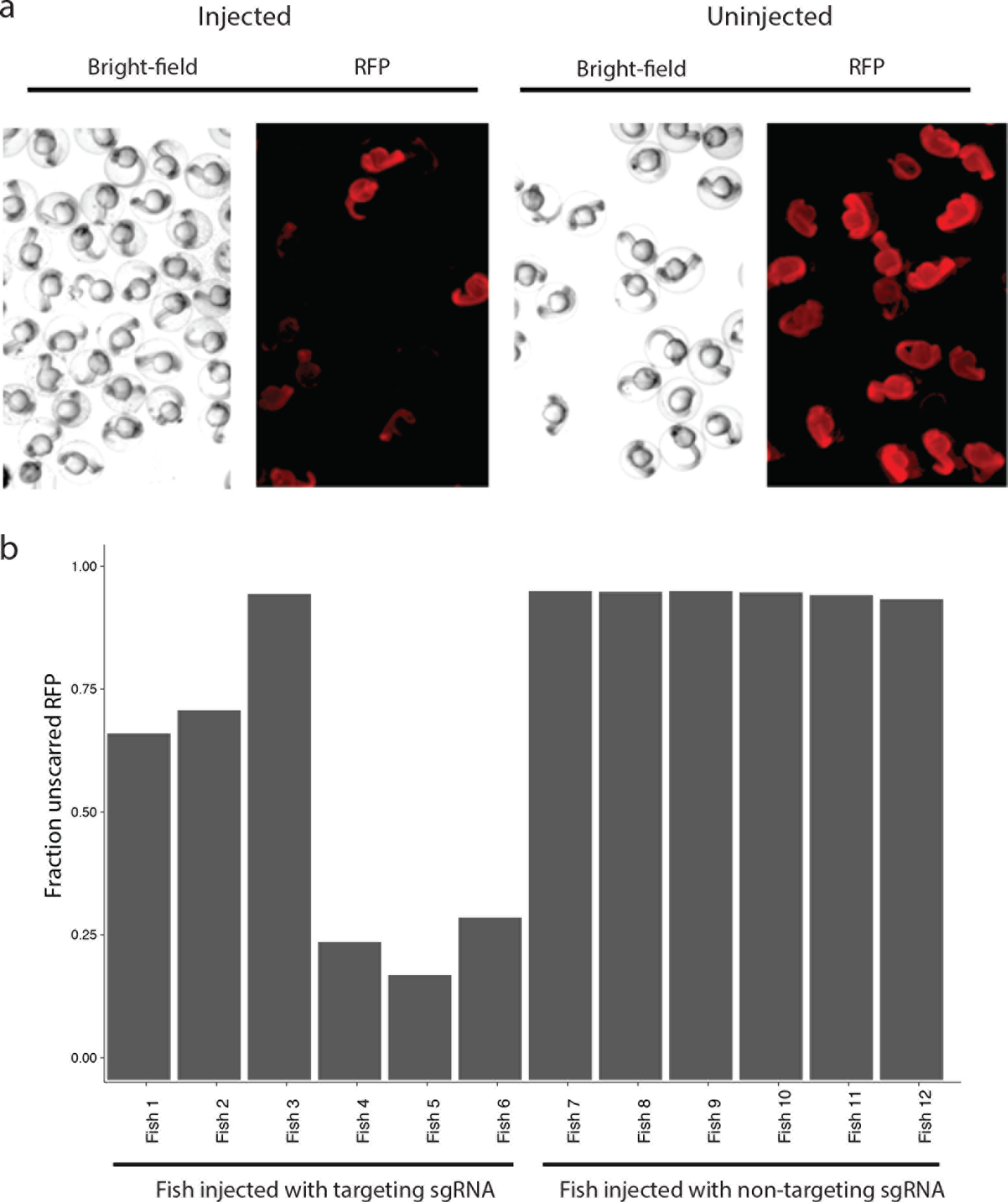
Detection of scar formation using microscopy and sequencing. **(a)** Loss of RFP fluorescence upon injection of Cas9 and sgRNA targeting RFP. **(b)** Reduction of unscarred RFP upon injection of Cas9 and sgRNA targeting RFP.

**Supplementary Figure 2.**
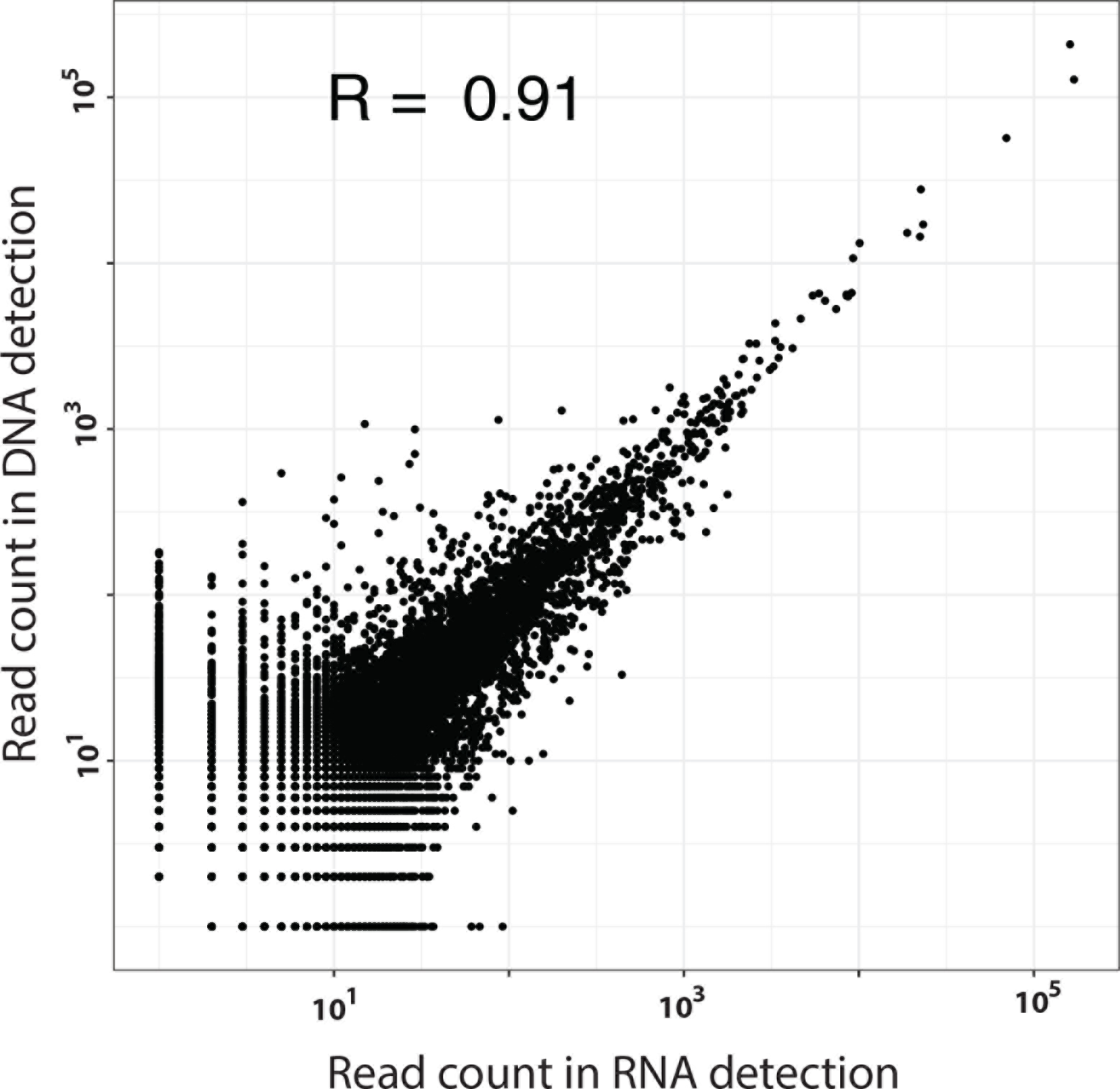
Strong correlation between DNA and RNA detection in the same embryo. Scar abundances detected in bulk on the DNA and RNA level for 5 dpf zebrafish larva.

**Supplementary Figure 3.**
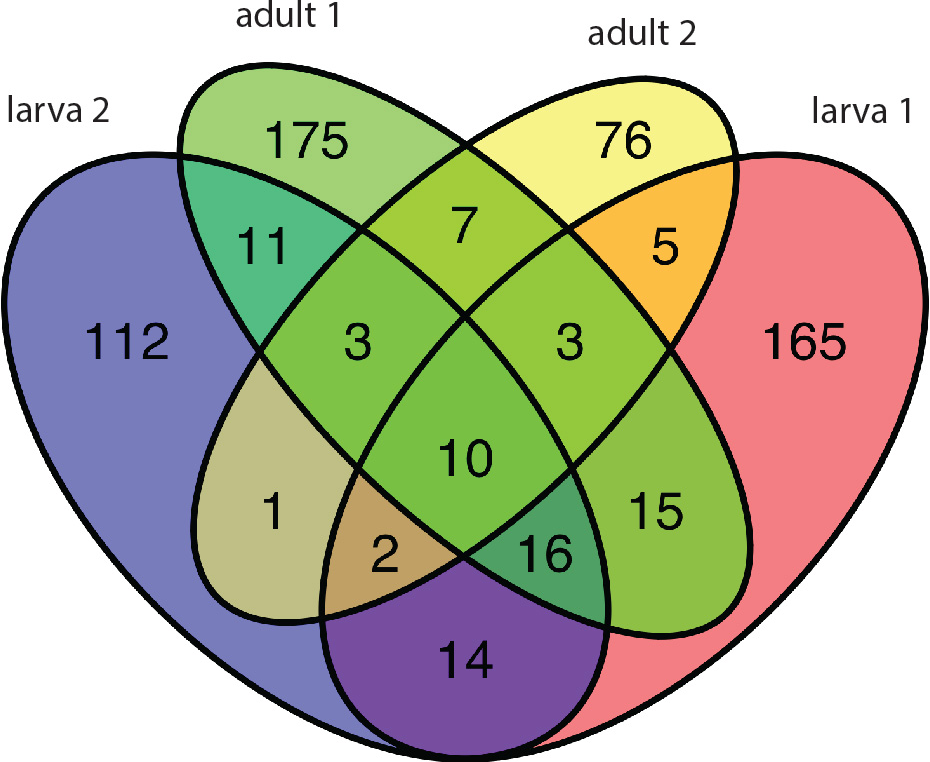
Scar diversity. Venn diagrams indicating numbers of unique and overlapping scars in four fish. “Larva 1” and “Adult 1” refer to the fish analyzed in the main figures, while “Larva 2” and “Adult 2” refer to the fish analyzed in replicate experiments.

**Supplementary Figure 4.**
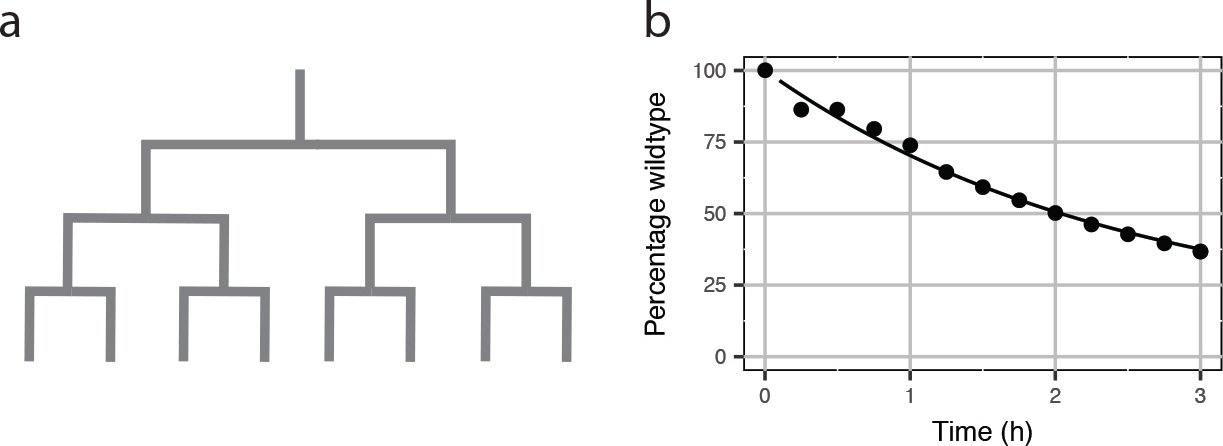
Scar probability modeling allows measurement of scarring rate. **(a)** The model requires two parameters, the previously measured cell division rate k_div_, and the per-site scarring rate k_scar_. We found that k_scar_ ≈ 0.3 /h fits the data best. **(b)** Scar creation dynamics resulting from these parameters.

**Supplementary Figure 5.**
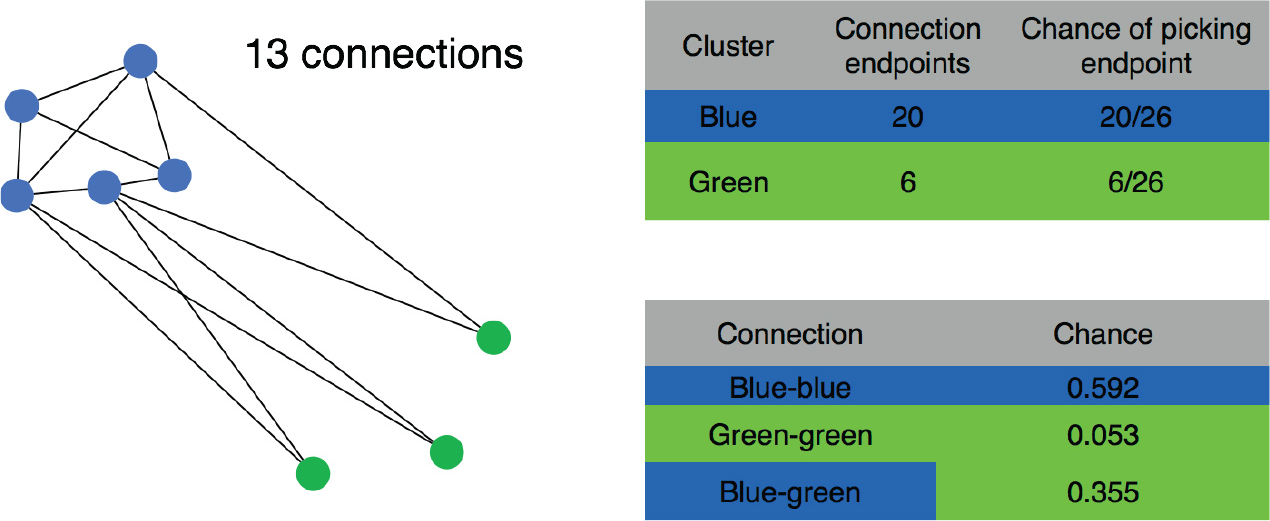
Example of background model for analysis of enriched connections. Given twenty connection endpoints in blue cells, and six in green cells, the chance of a blue endpoint is 10/13, and the chance of a green endpoint is 3/13. We can then calculate the chances of randomly selecting a blue-blue connection, a blue-green connection and a green-green connection. These chances determine a binomial distribution of connections to compare the observed connections against.

**Supplementary Figure 6.**
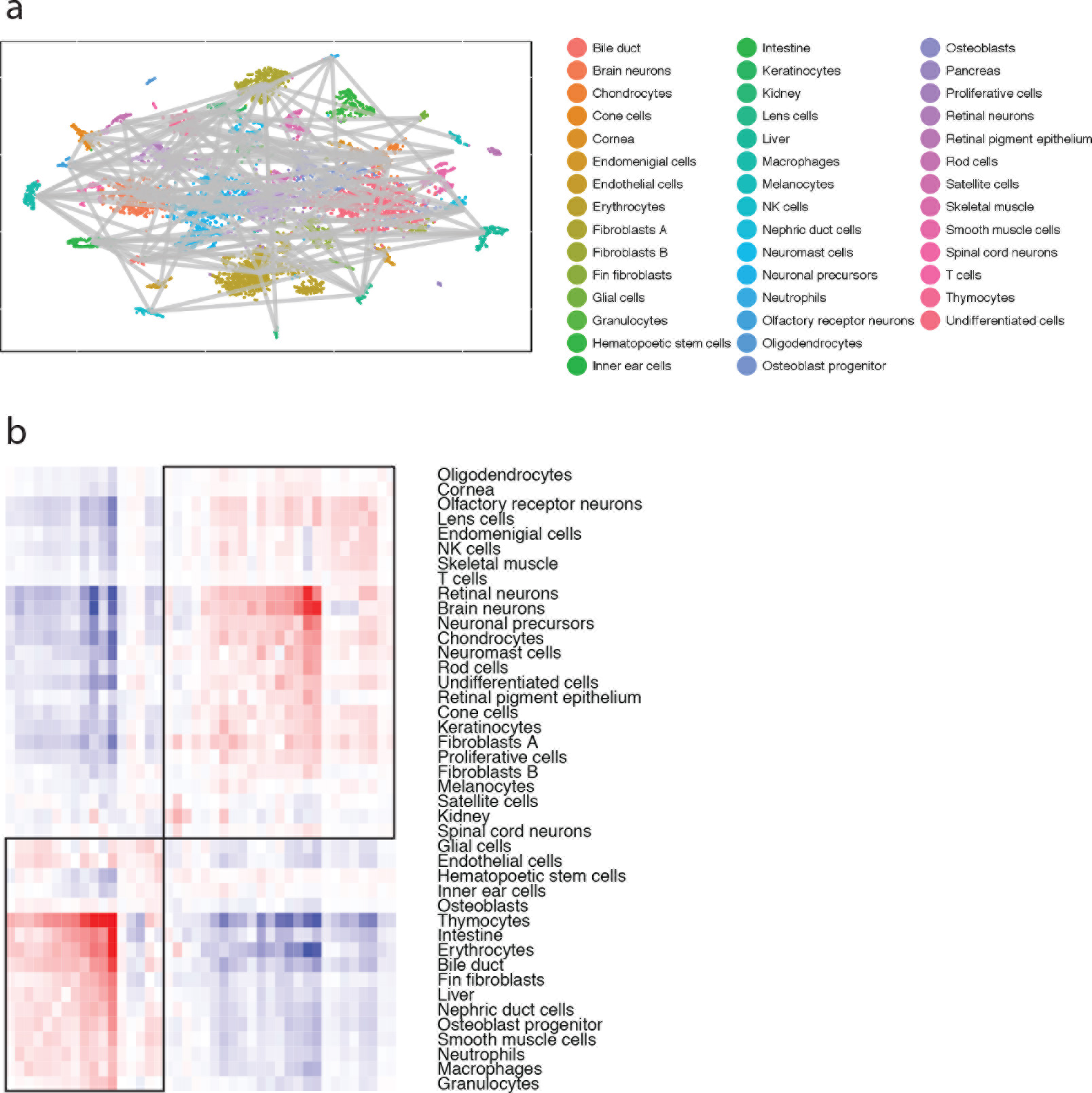
Enriched connections in another 5 dpf zebrafish larva. **(a)** Enrichments of scar connections between cell types compared to random distributions. **(b)** Hierarchical clustering of cell types by scar connection strength. See also Methods.

**Supplementary Figure 7.**
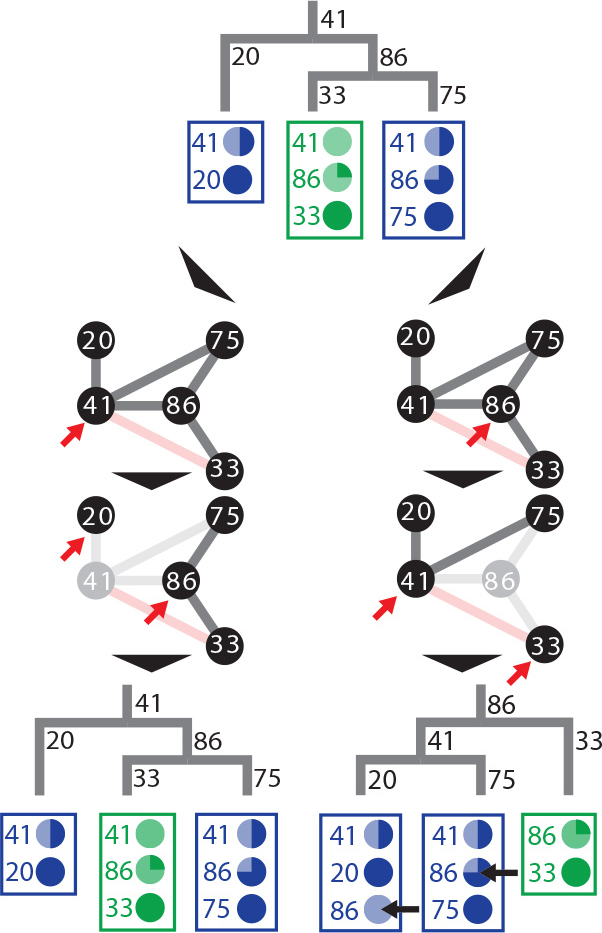
Evaluation and correction scheme for tree building algorithm. Summary cartoon of our computational approach for dealing with missing links. In this example, we start with a simple lineage tree that generates two cell types, “blue” and “green”, and we assume that scar 33, while being present, is not observed in any of the green cells. This leads to loss of a connection in the scar network graph (marked in pink), which in turn leads to ambiguity about which scar was created first. To evaluate the two reconstructed trees, we analyze the scar dropout rates in the resulting trees in a cell-type dependent manner. The tree on the right-hand side is unlikely, since it would require very different dropout rates for scar 86 in the two branches generating cell type “blue” (black arrows). The full evaluation and correction scheme is described in the Methods.

**Supplementary Figure 8.**
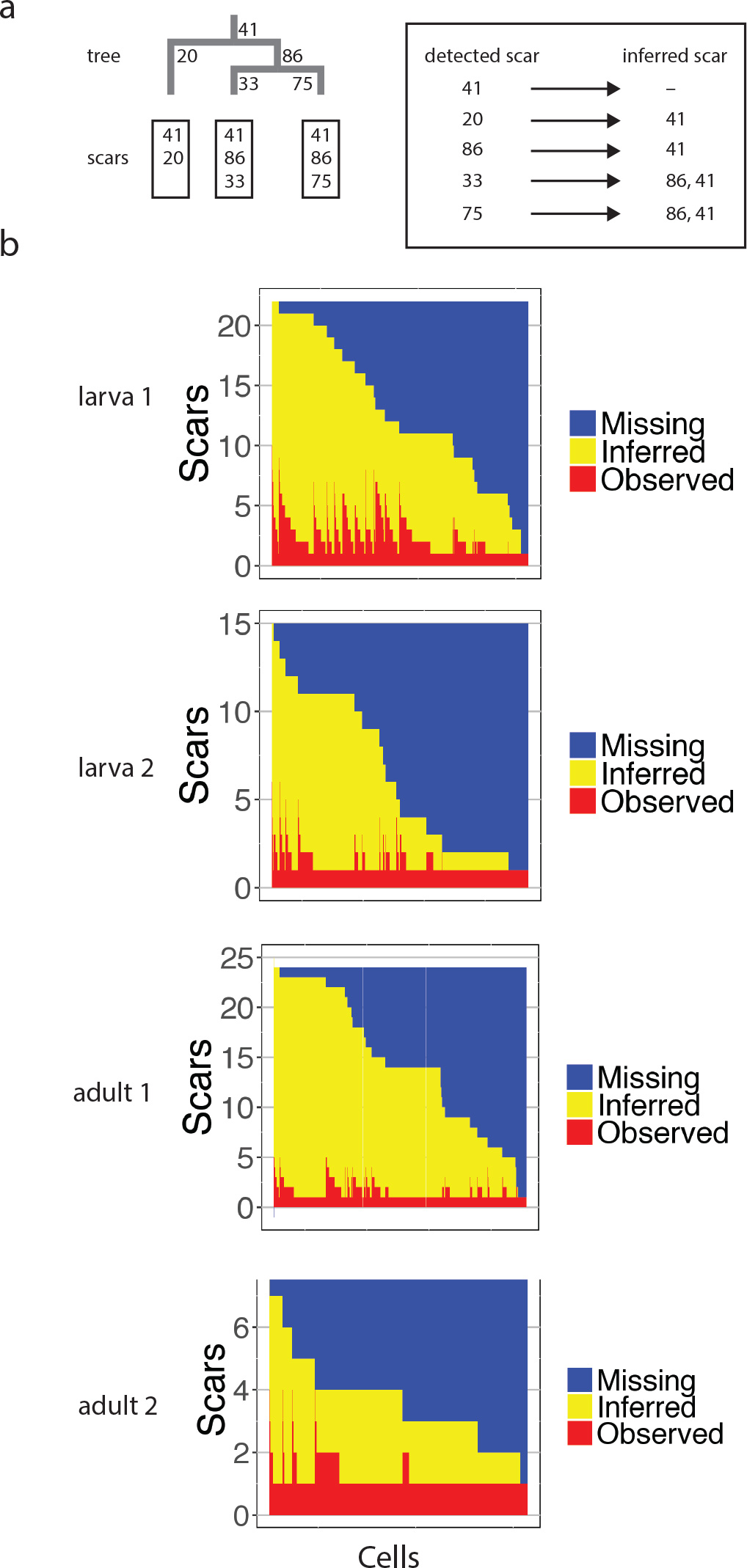
Lineage tree reconstruction allows inference of scars that were present but not detected. **(a)** Scar inference is based on the principle that, if the order of scarring is known, detection of a scar allows inference of all previously created scars. For instance, detection of scar 417 is sufficient for exact placement of a cell in the lineage tree in Fig. 2e, whereas detection of scar 312 without measurement of scars 260 and 417 only provides lineage information down to an intermediate level in the lineage tree. **(b)** Detected (red) and inferred (yellow) scars in single cell data.

**Supplementary Figure 9.**
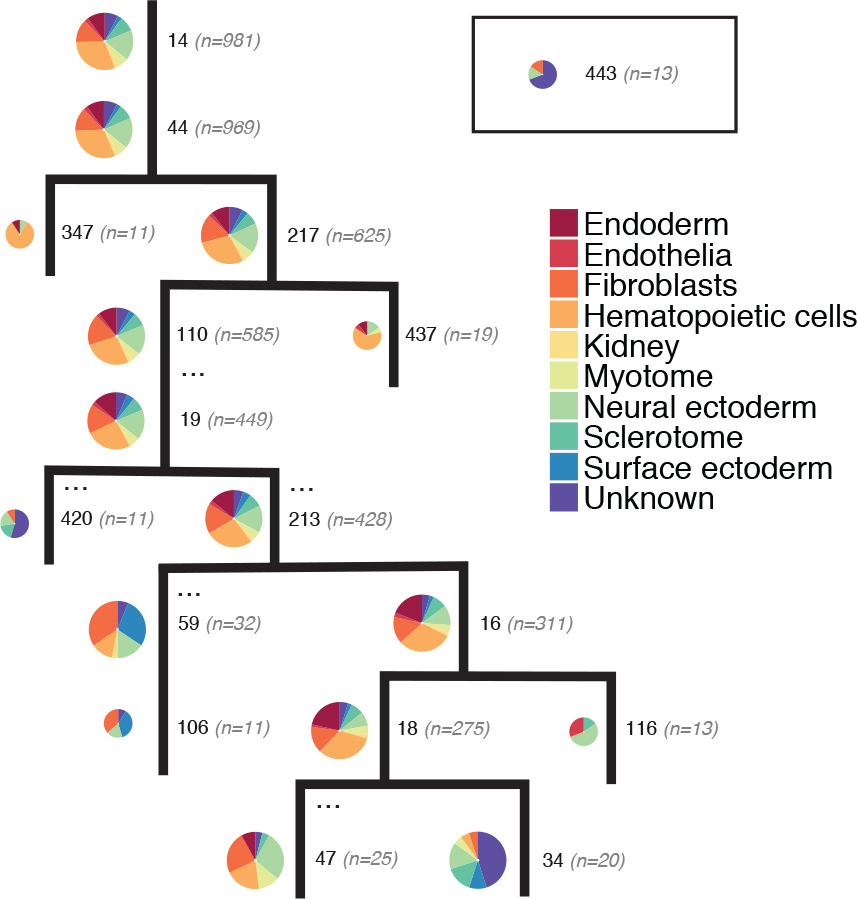
Lineage tree for another 5 dpf zebrafish larva. Lineage tree reconstruction and grouping of cell types as in Fig. 2e.

**Supplementary Figure 10.**
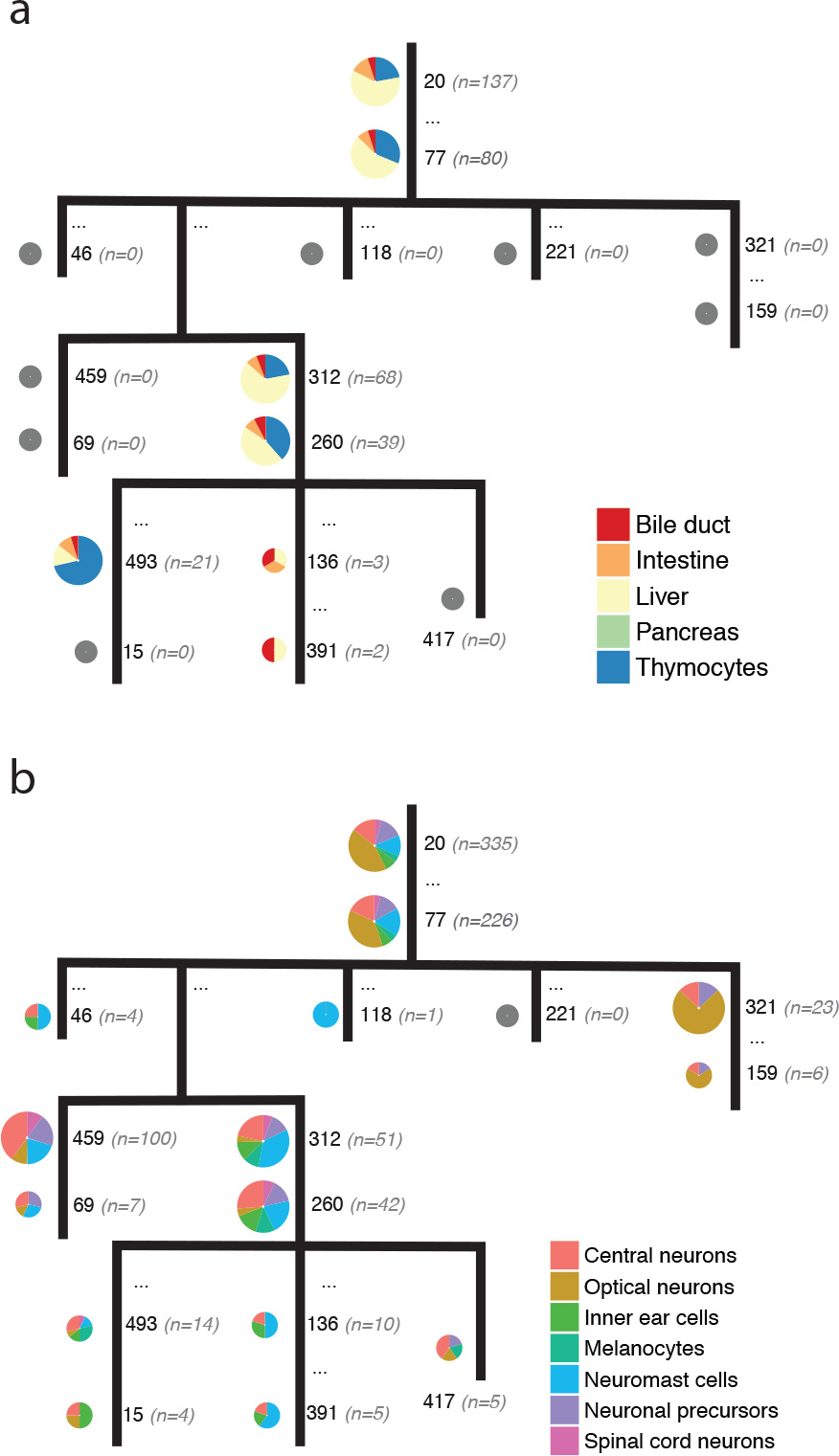
Endodermal and neuronal lineage trees in 5 dpf larva. Lineage trees for endodermal **(a)** and neuronal **(b)** cell types at higher cell type resolution for the 5 dpf larva analyzed in Fig. 2e, f. We observe cell-type specific lineage branches that give rise to different organs, such as the thymus, the hepatopancreas, and the optic apparatus.

**Supplementary Figure 11.**
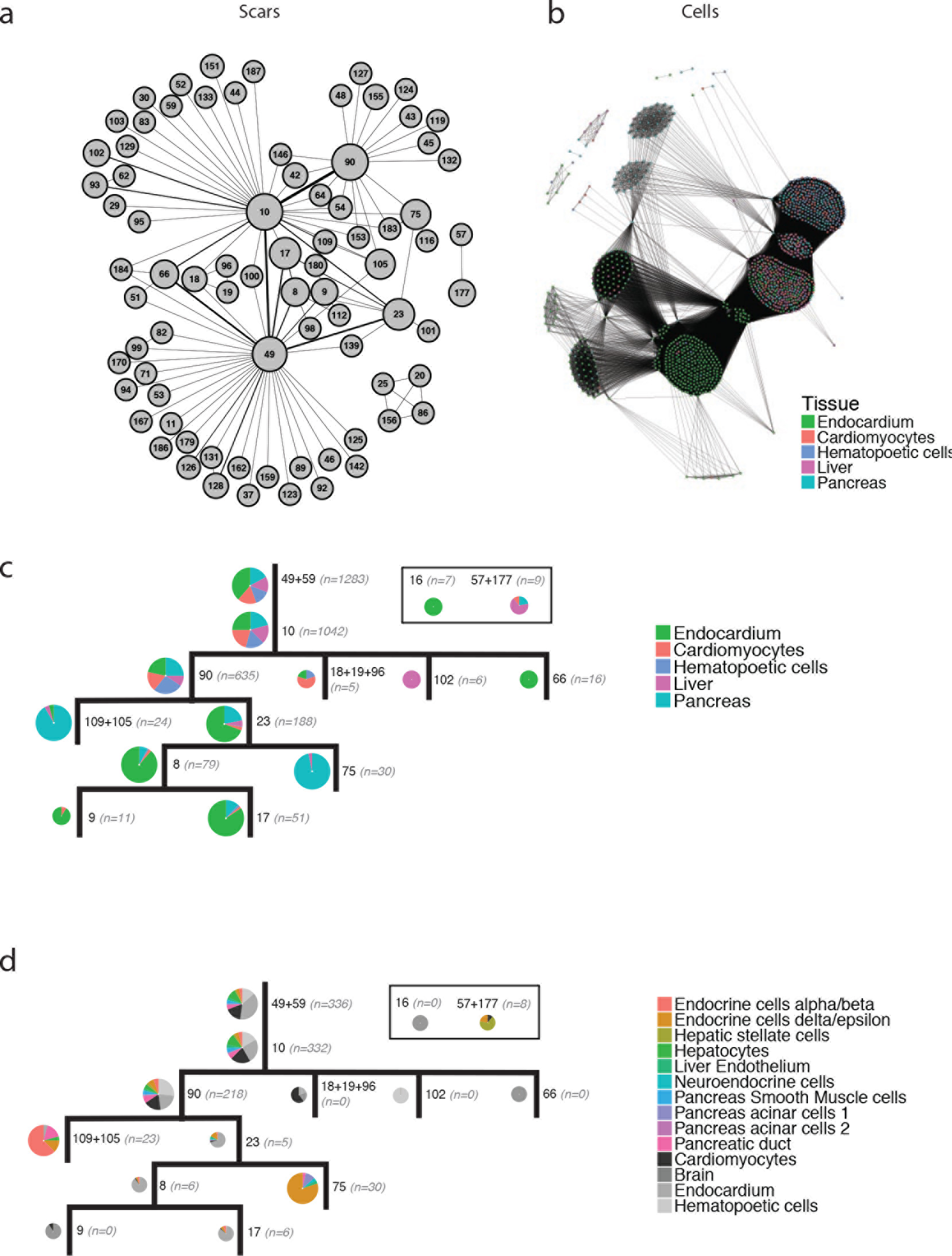
Replicate experiment for dissected organs of adult zebrafish. **(a, b)** Scar and cell network graphs. **(c)** Complete lineage tree. **(d)** Lineage tree zoomed into endodermal cell types, revealing a hierarchy of lineage splits in the hepatopancreatic system, including a separation of alpha/beta and delta/epsilon cells.

**Supplementary Figure 12.**
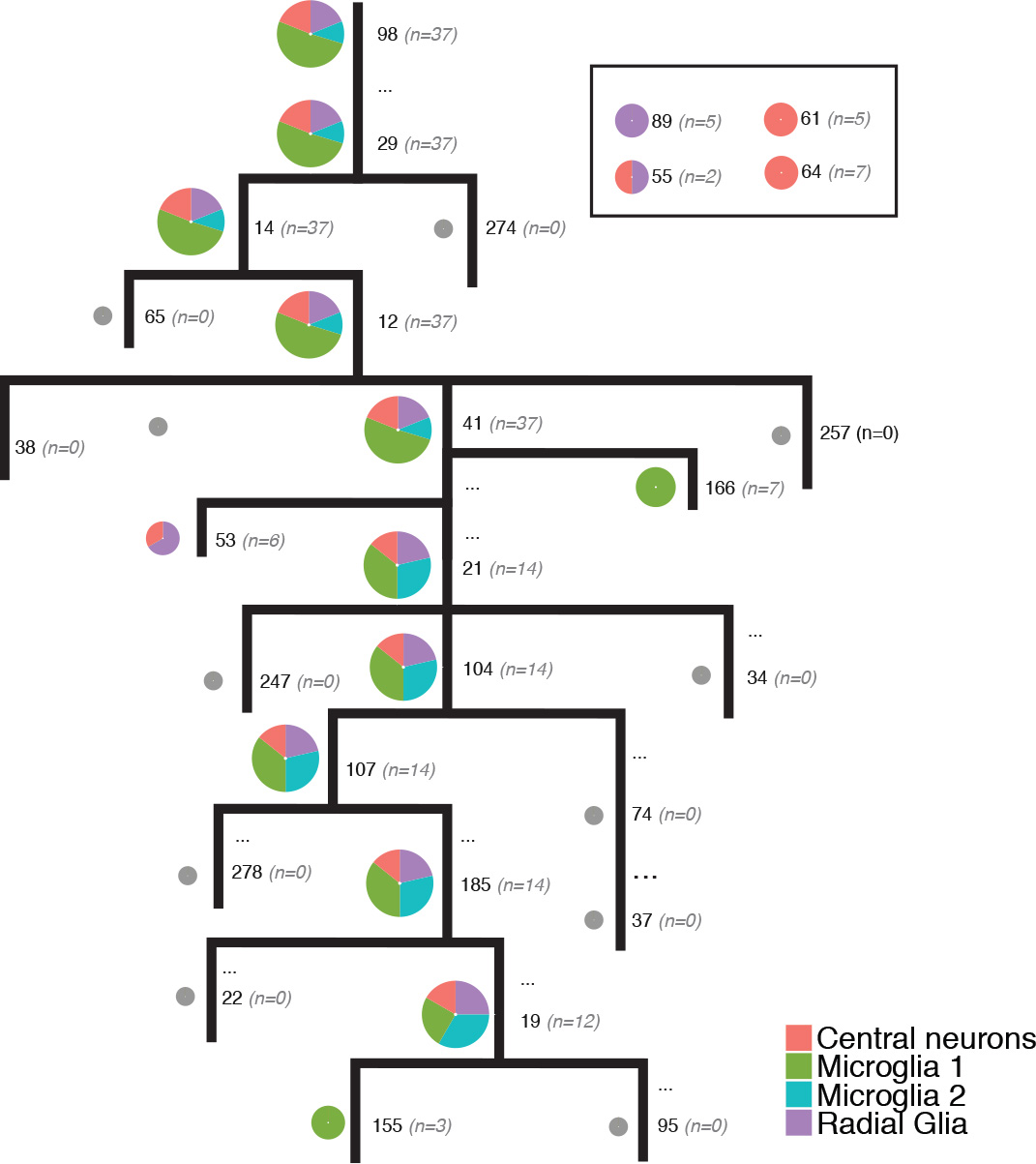
Neuronal cells in adult organs. Similar to Fig. 3e and 3f, we highlighted the cells of the neuronal lineage in analyzed adult organs. Most central nervous system neurons and radial glia are in clones separate from the main lineage tree, indicating an early precursor split. The three microglia with scar 155 share that scar with other immune cells.

